# Proteomic characterisation of Sarculator nomogram-defined risk groups in soft tissue sarcomas of the extremities and trunk wall

**DOI:** 10.1101/2023.11.14.567122

**Authors:** Madhumeeta Chadha, Sara Iadecola, Andrew Jenks, Jessica Burns, Amani Arthur, Valeriya Pankova, Christopher P Wilding, Dario Callegaro, Dirk C Strauss, Khin Thway, Alessandro Gronchi, Robin L Jones, Rosalba Miceli, Sandro Pasquali, Paul H Huang

## Abstract

**Background:** High-risk soft tissue sarcomas of the extremities and trunk wall (eSTS), as defined by the Sarculator nomogram, are more likely to benefit from (neo)adjuvant anthracycline-based therapy compared to low/intermediate-risk patients. The biology underpinning these differential treatment outcomes remain unknown.

**Methods:** We analysed proteomic profiles and clinical outcomes of 123 eSTS patients. A Cox model for overall survival including the Sarculator was fitted to individual data to define 4 risk groups. A DNA replication protein signature - Sarcoma Proteomic Module 6 (SPM6) was evaluated for association with clinicopathological factors and risk groups. SPM6 was added as a covariate together with Sarculator in a multivariable Cox model to assess improvement in prognostic risk stratification.

**Results:** DNA replication and cell cycle proteins were upregulated in high risk versus very low risk patients. Evaluation of the functional effects of CRISPR-Cas9 gene knockdown of proteins enriched in high risk patients identified candidate drug targets. SPM6 was significantly associated with tumour malignancy grade (p = 1.6e-06), histology (p = 1.4e-05) and risk groups (p = 2.6e-06). Cox model analysis showed that SPM6 substantially contributed to a better calibration of the Sarculator nomogram (Index of Prediction Accuracy =0.109 for Sarculator alone versus 0.165 for Sarculator + SPM6).

**Conclusions:** Risk stratification of patient with STS is defined by distinct biological pathways across a range of cancer hallmarks. Incorporation of SPM6 protein signature improves prognostic risk stratification of the Sarculator nomogram. This study highlights the utility of integrating protein signatures for the development of next-generation nomograms.

## Introduction

Around half of soft tissue sarcomas (STS) arise in the extremities or trunk wall and comprise a broad range of different histological types with wide variation in clinical outcomes^1^. Several prognostic nomograms based on baseline clinical and pathological variables have been developed to predict survival outcomes after surgical resection of primary localised STS with curative intent, which can aid clinical management^2^. Of these nomograms, the predictive accuracy of the Sarculator tool has been independently validated in multiple large international series^3,4^. More recently, retrospective analyses of two randomised trials have shown that patients with high risk STS of the extremities or trunk wall (as defined by the Sarculator nomogram) benefitted from peri-operative anthracycline-based therapy, while low/intermediate-risk patients did not^5,6^. These findings suggest that the nomogram-predicted risk groups may have distinct biology which could explain their differential responses to treatment. However, the biological features of these risk groups have yet to investigated. Furthermore, there is currently a lack of effective targeted agents beyond anthracyclines in the (neo)adjuvant setting to improve cure rates following surgical resection, and in-depth characterisation of the molecular pathways in tumours from high risk patients may lead to the discovery of new drug targets for use in this setting. In this respect, proteomic data may be particularly useful as unlike other biomolecules such as DNA and RNA, proteins represent the largest druggable class of oncology targets with direct translational applicability for drug discovery and repurposing^7,8^.

The Sarculator nomogram includes the covariates of baseline tumour size and malignancy grade, histological type and patient age^3^. While the predictive accuracy of this nomogram is relatively high, there remains an opportunity for further improvements by inclusion of other biological factors that capture orthogonal information beyond these baseline clinicopathological variables. Indeed, recent efforts in integrating an inflammatory biomarker prognostic index showed a significant improvement in the discriminative ability of the Sarculator in primary retroperitoneal sarcoma patients^9^. Along similar lines, two studies have recently evaluated the ability of the gene expression-based CINSARC signature to enhance the prognostic accuracy of Sarculator with contrasting results^10,11^. Integrating biological and molecular signatures with nomograms is an emerging field and holds the promise of next generation nomograms that capture both biological and clinical features for improved prognostication^2,12^.

Here we investigate the protein networks that are characteristic of nomogram-stratified high risk and low risk STS of the extremities or trunk wall and evaluate a specific protein signature Sarcoma Proteomic Module 6 (SPM6) as a complementary prognostic tool to the Sarculator nomogram.

## Methods and material

### Patient cohort

The cohort is comprised of 123 patients with primary STS of the extremities or trunk wall from The Royal Marsden Hospital. Retrospective collection and analysis of associated clinical data was approved as part of the Royal Marsden Hospital (RMH) PROgnoStic and PrEdiCTive ImmUnoprofiling of Sarcomas (PROSPECTUS) study (NHS Research Ethics Committee Reference 16/EE/0213). Baseline clinicopathological characteristics and survival data were collected by retrospective review of medical records^13^.

### Proteomic data

Proteomic data for this study were downloaded from ProteomeXchange (PXD036226) https://www.ebi.ac.uk/pride/archive/projects/PXD036226^13^. Raw mass spectra was processed using the SequestHT search engine in Proteome Discoverer 2.2 or 2.3 (Thermo Scientific, Waltham, MA, USA) and reviewed against UniProt human protein entries (v2018_07 or later). Precursor mass tolerance of 20 ppm and a fragment ion mass tolerance of 0.02 Da were used to identify tryptic peptides with a maximum of 2 missed cleavages. Fixed modifications of TMT6plex at N-terminus/lysine and Carbamidomethyl at cysteine, along with dynamic modifications of oxidation of methionine and deamidation of asparagine/glutamine were used in the search parameters. For estimating peptide confidence, alongside Percolator node, a Peptide False Discovery Rate (FDR) of 0.01 was employed with validation based on q-value and decoy database search. For protein quantification, the reporter ion quantifier node was applied with an integration window tolerance of 15 ppm. Integration method was based on the most confident centroid peak at the MS3 level, and only unique peptides were used for quantification, considering protein groups for peptide uniqueness. Additionally, peptide quantification required an average reporter signal-to-noise ratio greater than 3 to ensure the reliability of the quantified proteins. Proteins with an FDR < 0.01 and at least two identified peptides were used for subsequent analysis.

All data was processed using custom R scripts v.4.1.1 or later. Proteins identified in <75% of the samples were removed. Data normalisation and removal of batch effects was done by dividing the TMT intensities by the reference followed by imputation using k-nearest neighbour algorithm (k-NN)^14^. Values were log2-transformed followed by median centring across the samples and standardisation within the samples. To visualise the proteomics dataset, supervised clustering was performed using Pearson correlation distance. SPM6 has previously been reported in Burns et al.,^13^ and is calculated based on the median expression levels of 41 DNA replication proteins for each patient.

### Proteomics statistical analysis

All statistical tests were two-sided and where required, p values were adjusted to false discovery rate (FDR) using the Benjamini–Hochberg procedure to account for multiple comparisons^15^. Unless otherwise specified, analysis was performed using custom R scripts in R v4.1.1 or later.

To identify upregulated proteins in high-risk group and very low risk group, Wilcoxon-Mann-Whitney test was performed. Volcano plot was generated using EnhancedVolcano^16^ in R. Overrepresentation analysis was performed with ClusterProfiler in R^17^ using Hallmark gene sets^18,19^. Differentially expressed proteins were ordered by log2-fold change and proteins present in the dataset used as the background for overrepresentation analysis.

### CRISPR-Cas9 functional genomic data

Genome-wide CRISPR-Cas9 screening data was downloaded from the Cancer Cell Line Encyclopaedia (CCLE) portal (https://sites.broadinstitute.org/ccle)^20^. The CRISPRGeneEffect dataset (DepMap Public 22Q2) was used for analysis. Essential genes were previously determined in CCLE by calculating gene scores using the CHRONOS model^21^, which identified gene knockout fitness across the full CCLE dataset of all cell lines (gene score of 0 is equivalent to a gene which is not essential, whereas a score of ≤-1 indicates this as an essential gene). Using the model information file (https://depmap.org/portal/download/all/), STS cell lines were identified with the following histologies excluded, rhabdomyosarcoma, fibrosarcoma and undifferentiated pleomorphic sarcoma. For each STS cell line, gene scores were evaluated to determine whether loss of each gene resulted in cell death (gene score <-1) or reduced proliferation (gene score >-1 and <0).

### Nomogram statistical analysis

Binary association between SPM6 and categorical variables was analysed using the non-parametric Wilcoxon test or Kruskal-Wallis test, as appropriate, and represented graphically by stratified boxplots. The study endpoint was Overall Survival (OS). OS time was defined as the interval elapsing from surgery to death from any cause. Time was censored at the last follow-up for patients still alive. The OS curves were estimated using the Kaplan-Meier method and compared using the log-rank test. The association between OS and the Sarculator nomogram or SPM6 was investigated by fitting univariable Cox regression models. Five-year OS Sarculator nomogram predicted probabilities (pr-OS) were extracted from the Cox model to identify four risk groups using quartiles as cut-offs.

We evaluated the prognostic improvement when adding SPM6 signature to the Sarculator nomogram in a multivariable Cox model. Bi-variable Cox models with interaction between SPM6 and, respectively, tumour size, malignancy grade, histological type and patient’s age (covariates included in the Sarculator nomogram), and the nomogram score were also fitted to determine whether the prognostic strength of SPM6 varied according to different covariates or nomogram values.

SPM6 was modelled using three-knots restricted cubic splines to obtain a flexible fit^22^. The discriminative ability of the Cox models was quantified using the Harrell *C* index^23^, while Index of Prediction Accuracy (IPA) allowed to evaluate both the discriminative ability and the calibration of the models (the higher the better)^24^. Model performance was also assessed through calibration plots comparing the observed Kaplan-Meier 5-year OS probabilities with those predicted by the model. The median follow-up was estimated with the reverse Kaplan-Meier method^25^. The software used for the analysis is R Version 4.2.1. We considered a statistical test as significant when the corresponding p value was less than 5%.

## Results

### Patient cohort and Sarculator-based risk stratification

Our cohort is comprised of 123 patients with localised STS of the extremities or trunk wall across 8 histological types; baseline clinicopathological features are summarized in Table 1. There were no grade 1 tumours in this cohort. The median follow-up time was 72.5 (interquartile range 62.9-106.7) months and the 5-year OS probability was 50.3% (95% confidence interval: 41.6-60.8%). pr-OS as defined by Sarculator had a median value of 52% (interquartile range 39-69%). Applying cut-offs based on pr-OS quartiles, we identify four categories corresponding to high risk (pr-OS≤39%), intermediate risk (39%<pr-OS≤52%), low risk (52%<pr-OS≤69%) and very low risk (pr-OS>69%) patients. The Kaplan–Meier survival curve according to pr-OS cut-offs showed a statistically significant difference between the risk groups (p<0.001) (Figure 1).

**Table 1.** Clinicopathological characteristics of the cohort.

|  | (n=123); n (%) |
| --- | --- |
| **Age (years)**<br>Median<br>IQR | 64.78<br>50.1-74.7 |
| **Sex**<br>Female<br>Male | 68 (55.3%)<br>55 (44.7%) |
| **Tumour depth**<br>Deep<br>Superficial | 93 (75.6%)<br>30 (24.4%) |
| **Surgical margin**<br>R0<br>R1<br>R2<br>Rx | 62 (50.4%)<br>58 (47.2%)<br>1 (0.8%)<br>2 (1.6%) |
| **Tumour size (mm)**<br>Median<br>IQR | 80<br>53.5-110 |
| **Histological subtype**<br>Angiosarcoma<br>Alveolar soft part sarcoma<br>Clear cell sarcoma<br>Dedifferentiated liposarcoma<br>Epithelioid sarcoma<br>Leiomyosarcoma<br>Synovial sarcoma<br>Undifferentiated pleomorphic sarcoma | 2 (1.6%)<br>1 (0.8%)<br>3 (2.4%)<br>4 (3.3%)<br>8 (6.5%)<br>33 (26.8%)<br>28 (22.8%)<br>44 (35.8%) |
| **FNCLCC grade**<br>II<br>III | 77 (37.4%)<br>46 (62.6%) |
| **Chemotherapy**<br>done<br>not done<br>NA | 11 (9.0%)<br>111 (90.2%)<br>1 (0.8%) |
| **Radiotherapy**<br>done<br>not done<br>NA | 10 (8.1%)<br>112 (91.1%)<br>1 (0.8%) |

**Figure 1.**
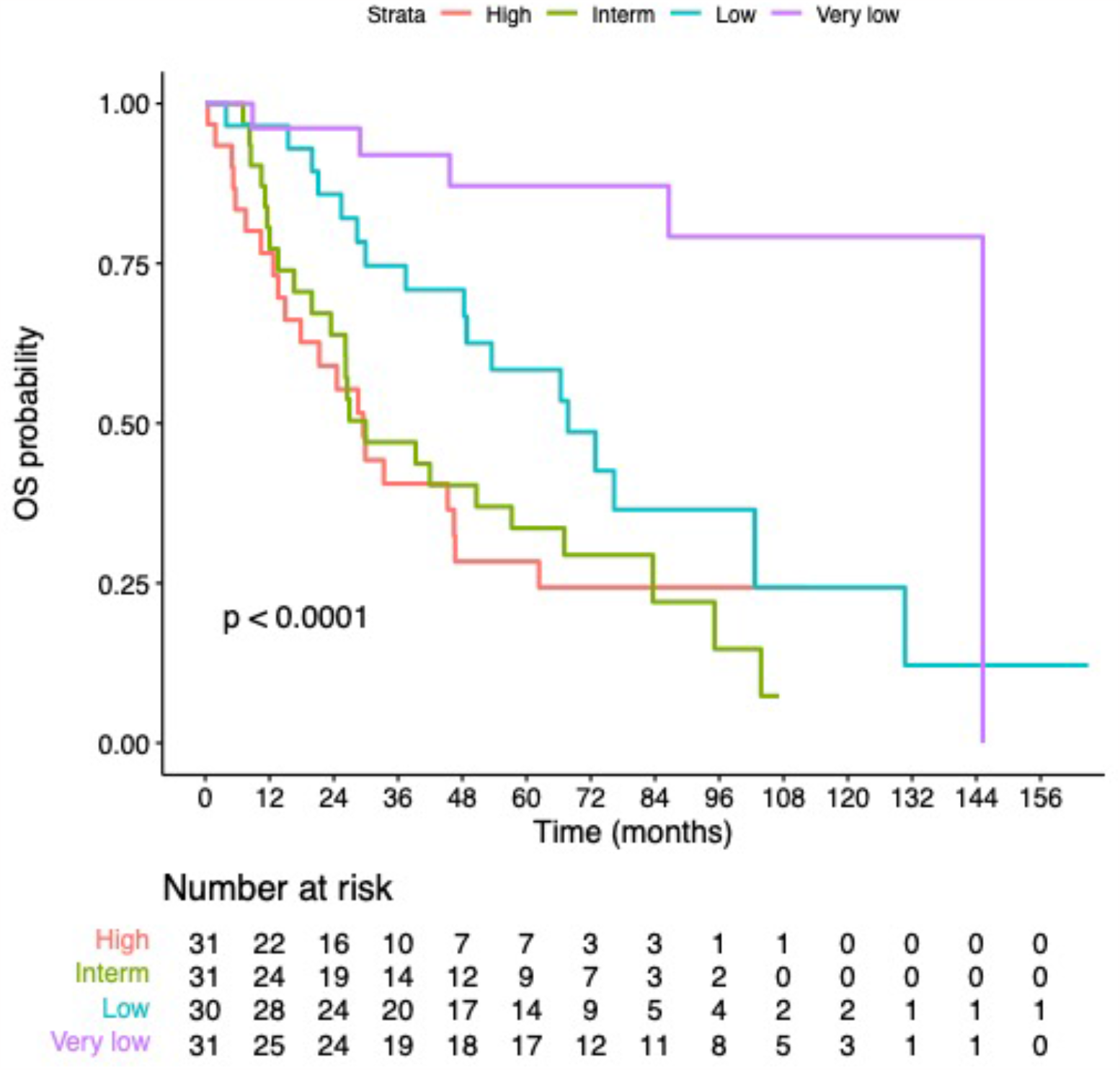
Kaplan-Meier curves for overall survival according to Sarculator predicted risk groups

### Proteomic features of nomogram predicted risk groups

We have undertaken mass spectrometry-based analysis to characterise the tumour proteomic profiles in this cohort^13^. Supervised clustering of the full proteomic dataset (n=3419 proteins, Table S1) of the 4 nomogram predicted risk groups is shown in Figure 2A. Proteins that were significantly enriched in the high risk versus the very low risk groups (n=62 patients and 3459 proteins) were identified by Wilcoxon-Mann-Whitney test with multiple testing correction (Figure 2B). This analysis identified 44 proteins that were significantly upregulated in the nomogram-predicted high risk group (adj p <0.05, fold change >2) and 44 proteins that were significantly upregulated in the nomogram predicted very low risk group (adj p<0.05, fold change >2) (Table S2). Proteins that were upregulated in the high risk group include components of the minichromosome maintenance (MCM) complex (MCM2, MCM3, MCM5, MCM6 and MCM7), the cell cycle protein CDK1, and proteins involved in collagen crosslinking and proline hydroxylation (PLOD1, PLOD2, PLOD3). Proteins that were enriched in the very low risk group include mitochondrial matrix proteins involved in oxidative phosphorylation (ATP5F1, ATP5C1, SUCLA1, NDUFA9) and proteins regulating fatty acid oxidation (ACADVL, ACADS and CRAT). Consistent with these results, overrepresentation analysis of Hallmark gene sets finds that the high risk group of patients are enriched in E2F targets and G2M checkpoint proteins (Figure 2C) while very low risk patients were enriched in fatty acid metabolism.

**Figure 2.**
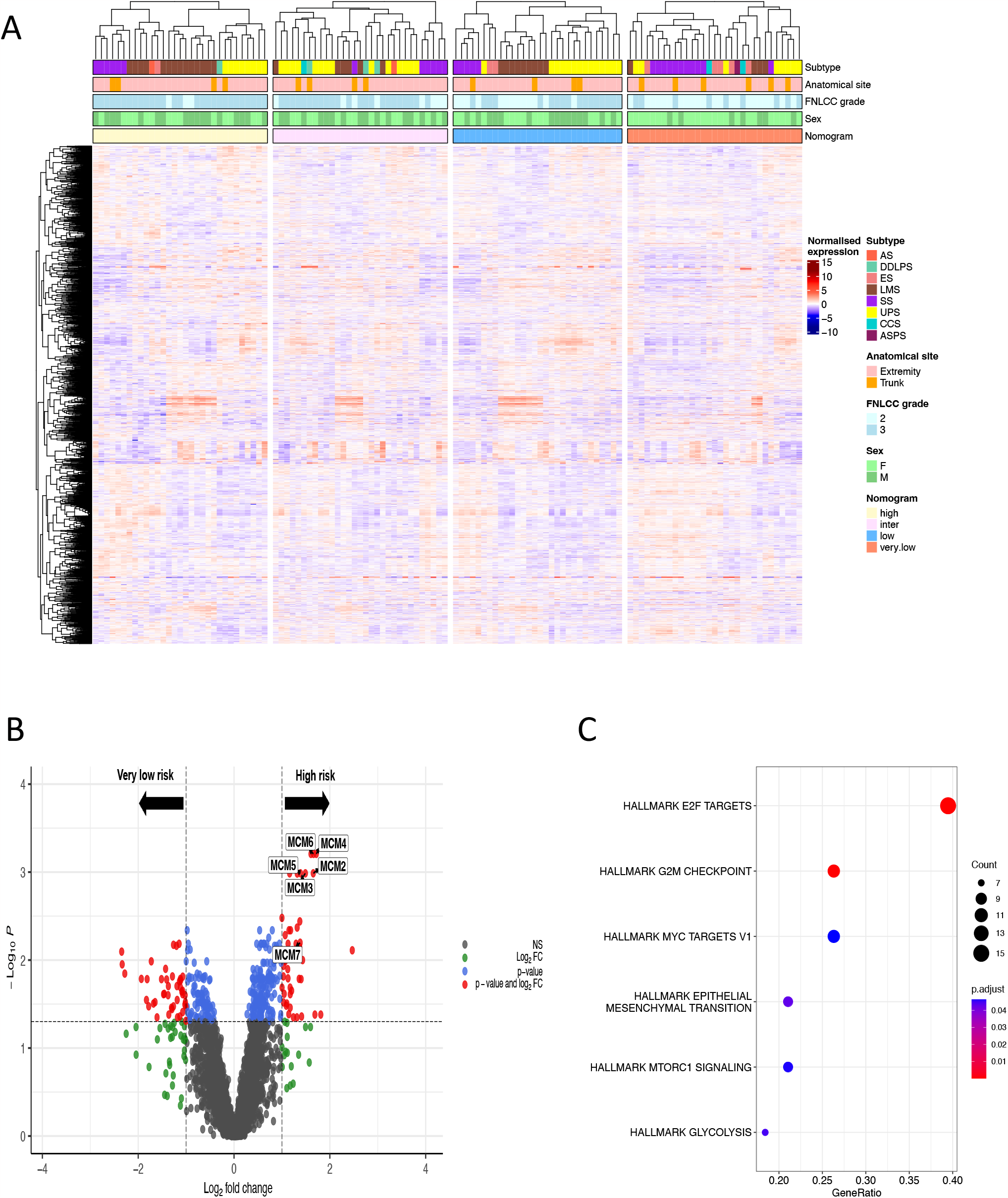
Proteomic analysis of nomogram predicted risk groups. (A) Annotated heatmap showing supervised clustering of the nanogram predicted risk groups for 3,419 proteins. Top to bottom panels indicate histological subtype, anatomical site, grade, sex and nomogram predicted groups. Abbreviations: AS = angiosarcoma; DDLPS = dedifferentiated liposarcoma; ES = epithelioid sarcoma; LMS = leiomyosarcoma; SS = synovial sarcoma; UPS = undifferentiated pleomorphic sarcoma; CCS = clear cell sarcoma; ASPS = alveolar soft part sarcoma. **(B)** Volcano plot showing significantly upregulated proteins in high risk and very low risk groups. Significant proteins (FDR<0.05, fold change >2) determined by multiple t-test followed by Benjamini Hochberg procedure are shown in red. MCM proteins significantly upregulated in the high risk group have been annotated. **(C)** Dot plot showing hallmark pathways overrepresented in the high risk group. The diameter indicates the number of proteins overlapping with Hallmark gene sets and colour indicates enrichment adjusted p values.

There is a need for new therapies to improve cure rates following surgery in high risk STS patients. To identify candidate drug targets from the list of 44 proteins that were significantly upregulated in nomogram predicted high risk patients, we evaluated the functional consequence of selective CRISPR-Cas9 knockdown of these genes in the cancer cell line encyclopaedia (CCLE) database^20^. We focused on 16 cell lines representing four histological types that are present in our proteomic dataset (leiomyosarcoma n = 4, liposarcoma n = 5, synovial sarcoma n = 5, epithelioid sarcoma n = 2) (Table S3). Genetic depletion of 11 genes (25%) caused cell death in at least 50% of the cell line panel. All these hits were known essential genes including the MCM complex, CDK1, PCNA and SRSF2 (Figure S1). In another 22 genes (50%), genetic depletion resulted in a decrease in cell viability in at least 50% of the cell line panel. Many of these genes are non-essential and include KPNA2, ENO1 and UAP1, which have previously been reported as drug targets in other cancer types (Figure S1)^26-29^.

### Evaluation of SPM6 signature in improving the prognostic risk stratification of the Sarculator nomogram

We have previously identified 14 SPM protein signatures which comprise a broad range of biological functions^13^. Given the enrichment of the MCM complex in nomogram predicted high risk patients, we evaluated if the SPM6 signature, a module comprising DNA replication proteins including components of the MCM complex (Table S4), had prognostic value in STS of the extremities or trunk wall. Median expression levels of SPM6 proteins for each patient was obtained^13^ and assessed for association of SPM6 with baseline clinicopathological variables of tumour grade and histological type. SPM6 was significantly associated with grade 3 tumours having higher levels of this variable compared to grade 2 cases (Wilcoxon p = 1.6e-06) (Figure 3A). In addition, SPM6 expression levels were associated with histological type with dedifferentiated liposarcoma cases having the lowest expression levels compared to angiosarcoma patients (Figure 3B). Furthermore, leiomyosarcoma patients had wide variation of SPM6 values indicative of broad heterogeneity of DNA replication protein expression levels within this histology. The global comparison of SPM6 levels between all histological types was statistically significant (Kruskal-Wallis p = 1.4e-05). We further evaluated the association of SPM6 protein expression levels with the four Sarculator predicted risk groups. There was a direct relationship between nomogram predicted risk groups and SPM6 with increasing risk being associated with increasing median SPM6 expression levels (Kruskal-Wallis p = 2.6e-06) (Figure 3C). In univariate Cox regression analysis, the Sarculator nomogram was significantly associated with patient OS (p<0.0001) while median SPM6 levels was not (p=0.242) (Table S5).

**Figure 3.**
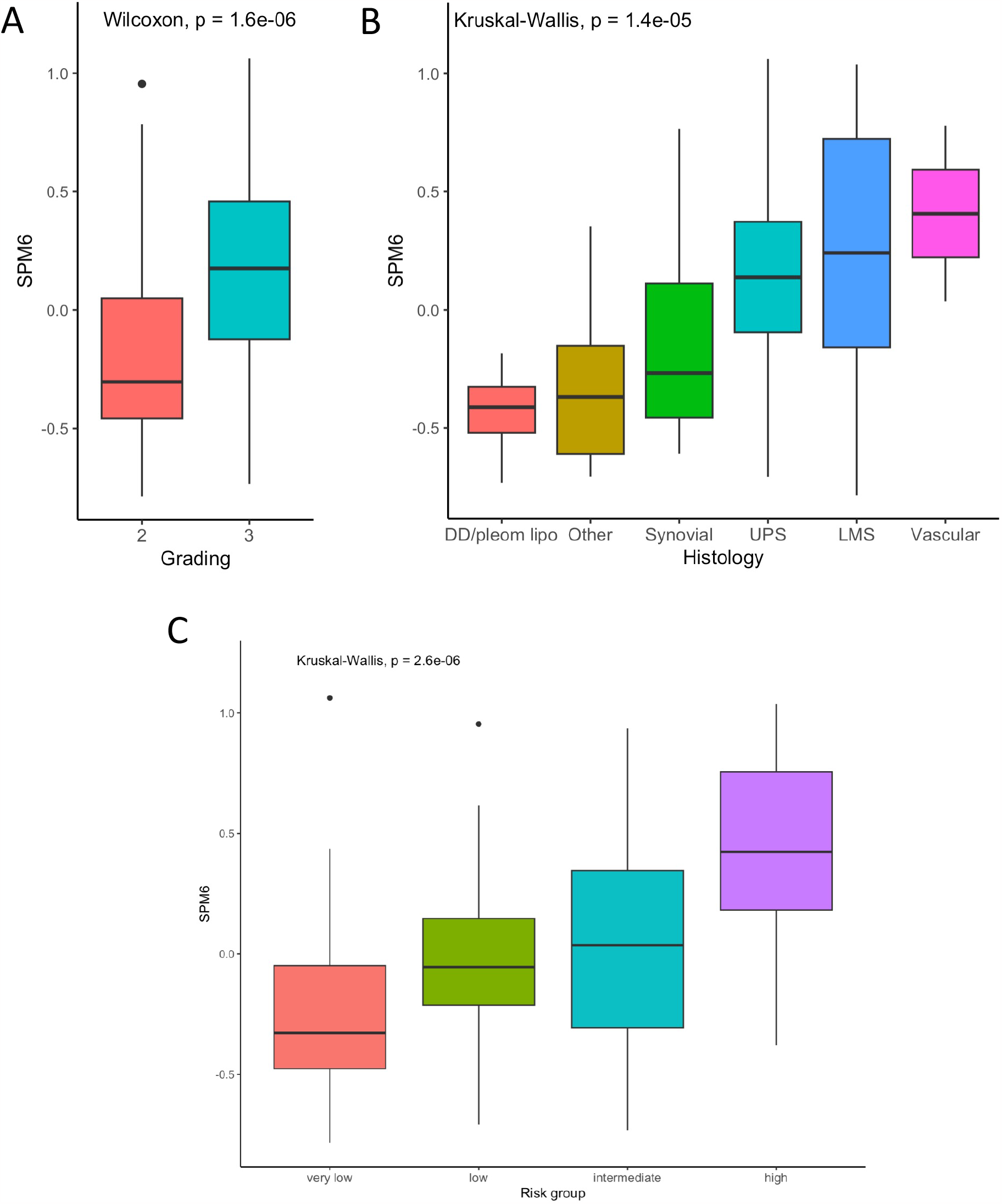
Association of SPM6 median expression protein levels with (A) tumour grade, (B) histology and (C) nomogram predicted risk groups.

Cox multivariable analysis showed that SPM6 slightly improved the discriminative ability of the Sarculator nomogram (Harrell C index=0.69 (95% confidence interval (CI) 0.62-0.759) for Sarculator nomogram alone vs 0.698 (95% CI 0.631-0.766) for Sarculator nomogram + SPM6). As regards to the calibration (Figure 4), while the nomogram predictions were quite accurate, despite the baseline survival recalibration operated in the present series, there was a discrepancy between predictions and observed outcomes particularly in the intermediate and very low risk groups, and addition of SPM6 substantially contributed to a better calibration (IPA = 0.109 vs 0.165).

**Figure 4.**
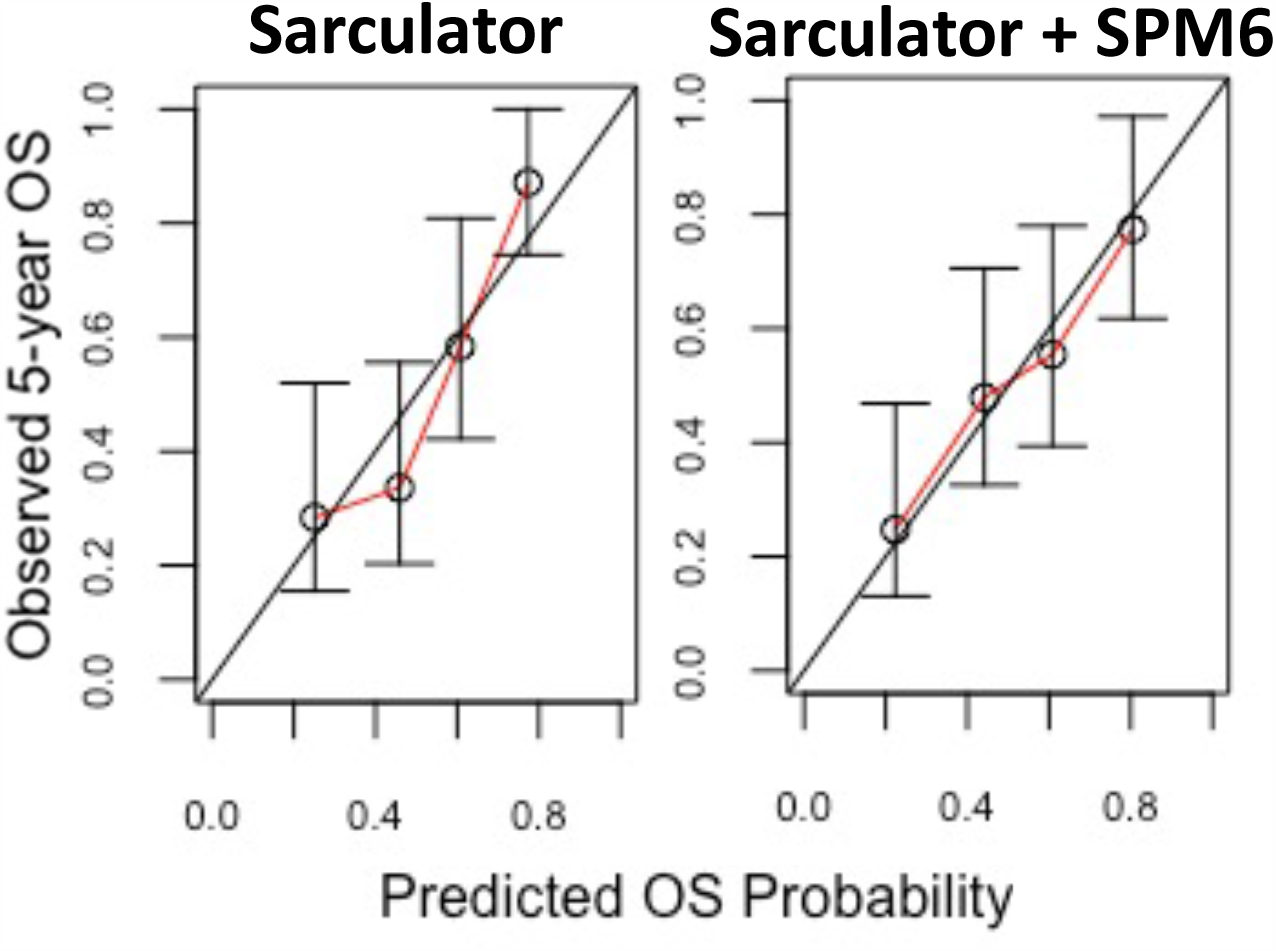
Calibration plots showing observed 5-year overall survival against predicted probabilities for Sarculator alone (left) and Sarculator + SPM6 (right).

In the additional analyses performed to determine whether the prognostic strength of SPM6 varied according to the nomogram or clinicopathological factors, we found that there was no significant interaction between the nomogram and SPM6 (Wald test SPM6 x nomogram p=0.705). Of the clinicopathological factors assessed (tumour malignancy grade, size, patient age and histological type), a statistically significant interaction was only detected between SPM6 and tumour size (Wald test SPM6 x size p=0.038) (Table S6) with the hazard ratios for SPM6 decreasing with increasing tumour size. This suggests that the SPM6 signature may provide additional orthogonal biological information particularly in the context of patients with smaller tumours and better OS outcomes.

## Discussion

In this study we characterised the proteins and biological pathways that underpin the prognostic risk stratification of extremities or trunk wall STS utilising the prognostic nomogram Sarculator. While the nomogram has been extensively used in clinical practice and evaluated in multiple clinical trial cohorts, the biology underpinning these predicted risk groups have hitherto been uncharacterised. Utilising high resolution mass spectrometry, we determine that high risk and very low risk patients are defined by distinct biological pathways across a broad range of cancer hallmarks including tumour cell proliferation, stromal microenvironment and metabolism^30^. Furthermore, we show that incorporation of a proteomic signature based on SPMs can improve the prognostic risk stratification of the Sarculator in extremity or trunk wall STS. To our knowledge, this is the first demonstration that inclusion of biological information in the form of proteomic signatures can improve on the prognostic accuracy of nomograms.

One of the main findings of this study is that high risk patients are characterized by enrichment in tumour proliferation and cell cycle regulation proteins belonging to the E2F targets and G2M checkpoint hallmark gene sets. For instance, high risk patients harboured an upregulation of the MCM complex which initiates DNA replication and is expressed prior to the G1 phase of the cell cycle^31^. An example is MCM3, a member of the MCM complex, has been shown to be associated with tumour cell proliferation in multiple cancer types and in functional studies^32^. Proliferation in STS is mainly described by FNCLCC tumour malignancy grade which scores different histopathological tumour features, including necrosis, differentiation and mitotic count^33^. While Ki-67, a marker of disease proliferation in several solid tumours, has previously been assessed in STS to show a partial association between tumour proliferation and malignancy grade^34-38^, the molecular characteristics underpinning tumour malignancy grade in STS remains largely unknown. The enrichment of proteins that regulate DNA replication and the cell cycle including the MCM complex and CDK1 identified in this study adds new biological pathway information to the definition of high risk STS.

There is a gap in our current knowledge of the biology underlying anthracycline-based chemotherapy response in STS. Our results could provide a mechanistic explanation for the clinical observation that high risk STS patients are more likely to benefit from peri-operative anthracycline-based chemotherapy than low-risk patients. The stratification of patient risk with the Sarculator nomogram enabled a reanalysis of the EORTC-STBSG 62931 trial, which failed to show a survival benefit for adjuvant anthracycline plus ifosfamide in STS^39^ and was for a long time considered as evidence for the lack of efficacy of chemotherapy in this setting. However, when patients with STS of the extremities or trunk within this study were stratified according to their Sarculator-predicted risk of death, a benefit of chemotherapy was detected only in the high risk patients^6^. This is consistent with anthracycline being more effective in tumour cells characterized by high levels of proliferation^40-42^. The enrichment of E2F targets and G2M checkpoint proteins in high risk patients identified in our study may explain in part the predictive value of the Sarculator in identifying these patients who are likely to have a reduced risk of recurrence when peri-operative anthracycline therapy is used. Consistent with our findings, similar functional hallmark gene sets have been reported among predictive factors for complete response to neoadjuvant anthracycline in other solid tumours, such as triple negative breast cancer^43^.

While there is evidence that peri-operative chemotherapy may benefit high risk STS patients, there remains broad inter-patient heterogeneity in tumour responses within this group. Furthermore, some histologies such as epithelioid sarcomas and clear cell sarcomas are known to be resistant to conventional chemotherapy^44^. Here we leverage on the proteomic profile of high risk STS patients to interrogate genome-wide CRISPR-Cas9 functional screens of STS cell lines within the CCLE database. This led to the rational nomination of additional candidate drug targets beyond anthracycline-based therapy. Of the non-essential genes identified to have a functional effect in decreasing cell viability in >50% of the cell lines assessed, genetic knockdown of KPNA2 displayed one of the strongest effects. KPNA2 is a member of the karyopherin family of nuclear export proteins and has been shown to have prognostic value in multiple cancer types including breast and prostate cancer^45-48^. Furthermore, this protein regulates tumour growth and migration in preclinical models of several cancer types including liver and gallbladder cancer^49,50^. While there are currently no drugs that directly targets KPNA2, the deubiquinating enzyme USP1 has been shown to regulate the stabilisation of KPNA2^51^. Importantly inhibitors to USP1 such as pimozide (which has been clinically used for patients with schizophrenia^52^) or more selective compounds such as ML323^53^ are able to substantially reduce the expression of KPNA2 which diminished breast cancer metastasis in vivo^51^. While hypothesis generating, our approach of integrating proteomic and functional genomics data can propose new candidate drug targets which could be used as alternatives to or in combination with anthracycline-based chemotherapy. These candidates need to be experimentally tested in well characterised patient-derived preclinical models of high risk STS to validate their effectiveness.

These results also have implications for improving the performance of Sarculator. Calibration of Sarculator, that is the correlation between predicted and observed survival, was improved by the addition of the prognostic information encoded by the SPM6 signature in this series. This shows that, although there was a direct correlation between nomogram predicted risk groups and SPM6, the latter has the potential of refining the prognostic prediction of the nomogram. SPM6 is comprised of proteins that are predominantly involved in the regulation of DNA replication. It is therefore interesting that this protein signature was correlated with different tumour characteristics that describe disease biology, such as tumour malignancy grade and histology, but not with tumour size. We further find that SPM6 may add orthogonal biological information particularly in patients with smaller tumours and better OS outcomes. Although the inclusion of proteomic information is promising, this needs to be balanced with the availability of such molecular information in routine clinical practice. One of the major reasons for the widespread utility of Sarculator among clinicians is its use of easy-to-obtain and reproducible clinicopathological information that constitute the backbone of prognostic predictions. In contrast, proteomics is currently primarily a research use only tool which makes it challenging to implement in routine clinical management^7,8^. Whether the new prognostic information based on proteomic signatures such as SPM6 can be incorporated to the Sarculator in future will require independent validation and larger patient numbers.

This study is limited by its retrospective study design. Patient selection based on availability of tumour tissue for proteomic analysis may have introduced a selection bias. In addition, the relatively small number of patients did not allow for a deep analysis of the possible differences among STS histologies. The absence of grade I tumours characterizes this cohort as a relatively homogenous higher risk group of patients. Interestingly, our proteomic analysis showed that the very low risk patients harboured features of metabolic rewiring with an enrichment of proteins involved in oxidative phosphorylation and fatty acid oxidation. Future proteomic analysis of grade 1 tumours may identify additional pathways that will improve our mechanistic understanding of the biology of low-risk STS patients. Nevertheless, these findings should be considered as hypothesis-generating and future validation in independent cohorts as well as functional experiments are required.

## Acknowledgements

This study is funded by grants from the Sarah Burkeman Trust, The Institute of Cancer Research, and the National Institute for Health Research (NIHR) Biomedical Research Centre at The Royal Marsden NHS Foundation Trust and The Institute of Cancer Research, and a charitable donation from Geoff Crocker and Bristol Care Homes to P.H.H, AIRC Individual Grant – Next Gen Clinician Scientist (28546) and Fondazione Regionale per la Ricerca Biomedica (1751036) to S.P, and The International Accelerator Award funded by Cancer Research UK (C56167/A29363)/ AIRC (24297) / Fundacion Cientifica - Asociacion Espanola Contra el Cancer (Foundation AECC-GEACC19007MA). The funders of the study had no role in the study design, data collection, data analysis, data interpretation or in writing of the manuscript.

## Data Availability

The raw proteomic data use in this study have been deposited in the ProteomeXchange Consortium via the PRIDE partner repository^54,55^ with the dataset identifier PXD036226 [https://www.ebi.ac.uk/pride/archive/projects/PXD036226]. The clinical data is available under restricted access due to data privacy legislation, access can be obtained by contacting the corresponding author (P.H.H) and will require the researcher to sign a data access agreement with the Institute of Cancer Research after approval by the Data Access Committee (DAC).

## Competing Interests Statement

The authors declare no competing interests.

## Figure legends

**Figure S1.**
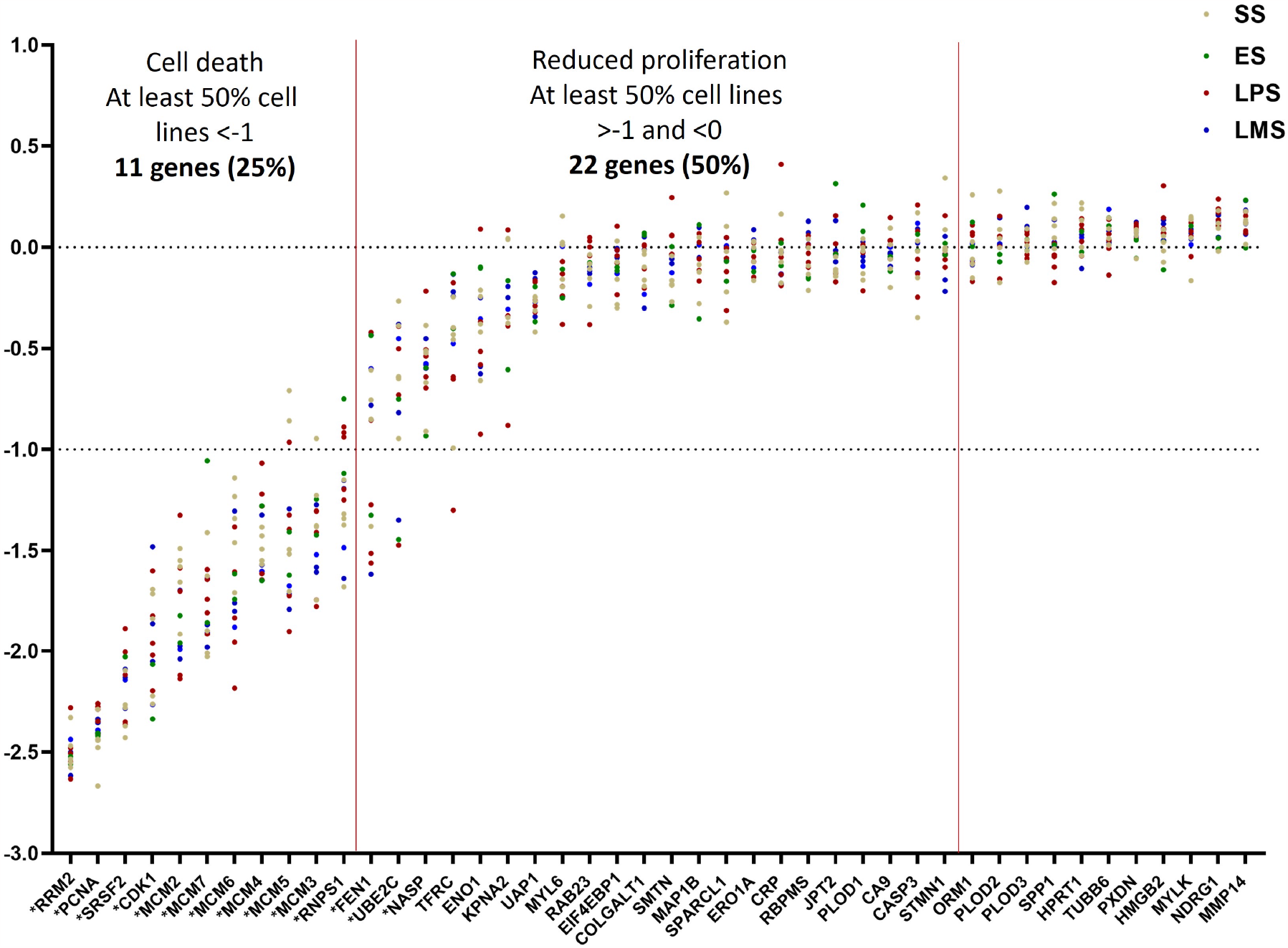
Analysis of CRISPR-Cas9 gene depletion data of 44 proteins enriched in high risk group in a panel of 16 STS cell line in the CCLE database (Table S3). * indicates essential genes. Abbreviations: SS = synovial sarcoma, ES = epithelioid sarcoma, LPS = liposarcoma and LMS = leiomyosarcoma.

## Notes

### Competing Interest Statement

The authors have declared no competing interest.

## References

1. Gronchi, A., et al. Soft tissue and visceral sarcomas: ESMO-EURACANGENTURIS Clinical Practice Guidelines for diagnosis, treatment and follow-up(☆). Ann Oncol 32, 1348–1365 (2021).

2. Danieli, M. & Gronchi, A. Staging Systems and Nomograms for Soft Tissue Sarcoma. Curr Oncol 30, 3648–3671 (2023).

3. Callegaro, D., et al. Development and external validation of two nomograms to predict overall survival and occurrence of distant metastases in adults after surgical resection of localised soft-tissue sarcomas of the extremities: a retrospective analysis. Lancet Oncol 17, 671–680 (2016).

4. Voss, R.K., et al. Sarculator is a Good Model to Predict Survival in Resected Extremity and Trunk Sarcomas in US Patients. Ann Surg Oncol (2022).

5. Pasquali, S., et al. Neoadjuvant chemotherapy in high-risk soft tissue sarcomas: A Sarculator-based risk stratification analysis of the ISG-STS 1001 randomized trial. Cancer 128, 85–93 (2022).

6. Pasquali, S., et al. The impact of chemotherapy on survival of patients with extremity and trunk wall soft tissue sarcoma: revisiting the results of the EORTC-STBSG 62931 randomised trial. Eur J Cancer 109, 51–60 (2019).

7. Burns, J., Wilding, C.P. R L.J. & P, H.H. Proteomic research in sarcomas - current status and future opportunities. Semin Cancer Biol 61, 56–70 (2020).

8. Chadha, M. & Huang, P.H. Proteomic and Metabolomic Profiling in Soft Tissue Sarcomas. Curr Treat Options Oncol 23, 78–88 (2022).

9. Fiore, M., et al. Preoperative Neutrophil-to-Lymphocyte Ratio and a New Inflammatory Biomarkers Prognostic Index for Primary Retroperitoneal Sarcomas: Retrospective Monocentric Study. Clin Cancer Res 29, 614–620 (2023).

10. Crombe, A., et al. Gene expression profiling improves prognostication by nomogram in patients with soft-tissue sarcomas. Cancer Commun (Lond) 42, 563–566 (2022).

11. Frezza, A.M., et al. CINSARC in high-risk soft tissue sarcoma patients treated with neoadjuvant chemotherapy: Results from the ISG-STS 1001 study. Cancer Med 12, 1350–1357 (2023).

12. Merry, E., Thway, K., Jones, R.L. & Huang, P.H. Predictive and prognostic transcriptomic biomarkers in soft tissue sarcomas. NPJ Precis Oncol 5, 17 (2021).

13. Burns, J., et al. The proteomic landscape of soft tissue sarcomas. Nat Commun 14, 3834 (2023).

14. Troyanskaya, O., et al. Missing value estimation methods for DNA microarrays. Bioinformatics 17, 520–525 (2001).

15. Benjamini, Y. & Hochberg, Y. Controlling the False Discovery Rate - a Practical and Powerful Approach to Multiple Testing. J R Stat Soc B 57, 289–300 (1995).

16. Blighe, K., Rana, S. & Lewis, M. EnhancedVolcano: Publication-ready volcano plots with enhanced colouring and labeling. (2023).

17. Yu, G., Wang, L.G., Han, Y. & He, Q.Y. clusterProfiler: an R package for comparing biological themes among gene clusters. OMICS 16, 284–287 (2012).

18. Liberzon, A., et al. The Molecular Signatures Database (MSigDB) hallmark gene set collection. Cell Syst 1, 417–425 (2015).

19. Liberzon, A., et al. Molecular signatures database (MSigDB) 3.0. Bioinformatics 27, 1739–1740 (2011).

20. Ghandi, M., et al. Next-generation characterization of the Cancer Cell Line Encyclopedia. Nature 569, 503–508 (2019).

21. Dempster, J.M., et al. Chronos: a cell population dynamics model of CRISPR experiments that improves inference of gene fitness effects. Genome Biol 22, 343 (2021).

22. Durrleman, S. & Simon, R. Flexible regression models with cubic splines. Stat Med 8, 551–561 (1989).

23. Harrell, F.E., Jr., Lee, K.L. & Mark, D.B. Multivariable prognostic models: issues in developing models, evaluating assumptions and adequacy, and measuring and reducing errors. Stat Med 15, 361–387 (1996).

24. Kattan, M.W. & Gerds, T.A. The index of prediction accuracy: an intuitive measure useful for evaluating risk prediction models. Diagn Progn Res 2, 7 (2018).

25. Schemper, M. & Smith, T.L. A note on quantifying follow-up in studies of failure time. Control Clin Trials 17, 343–346 (1996).

26. Han, Y. & Wang, X. The emerging roles of KPNA2 in cancer. Life Sci 241, 117140 (2020).

27. Huang, C.K., Sun, Y., Lv, L. & Ping, Y. ENO1 and Cancer. Mol Ther Oncolytics 24, 288–298 (2022).

28. Huang, L., et al. KPNA2 promotes cell proliferation and tumorigenicity in epithelial ovarian carcinoma through upregulation of c-Myc and downregulation of FOXO3a. Cell Death Dis 4, e745 (2013).

29. Yang, Y., et al. Discovery of Novel Inhibitors Targeting Multi-UDP-hexose Pyrophosphorylases as Anticancer Agents. Molecules 25(2020).

30. Hanahan, D. Hallmarks of Cancer: New Dimensions. Cancer Discov 12, 31–46 (2022).

31. Bochman, M.L. & Schwacha, A. The Mcm complex: unwinding the mechanism of a replicative helicase. Microbiol Mol Biol Rev 73, 652–683 (2009).

32. Ha, S.A., et al. Cancer-associated expression of minichromosome maintenance 3 gene in several human cancers and its involvement in tumorigenesis. Clin Cancer Res 10, 8386–8395 (2004).

33. Trojani, M., et al. Soft-tissue sarcomas of adults; study of pathological prognostic variables and definition of a histopathological grading system. Int J Cancer 33, 37–42 (1984).

34. Heslin, M.J., Cordon-Cardo, C., Lewis, J.J., Woodruff, J.M. & Brennan, M.F. Ki-67 detected by MIB-1 predicts distant metastasis and tumor mortality in primary, high grade extremity soft tissue sarcoma. Cancer 83, 490–497 (1998).

35. Huuhtanen, R.L., et al. Comparison of the Ki-67 score and S-phase fraction as prognostic variables in soft-tissue sarcoma. Br J Cancer 79, 945–951 (1999).

36. MacGrogan, G., et al. Comparison of quantitative and semiquantitative methods of assessing MIB-1 with the S-phase fraction in breast carcinoma. Mod Pathol 10, 769–776 (1997).

37. Rudolph, P., et al. Prognostic relevance of a novel proliferation marker, Ki-S11, for soft-tissue sarcoma. A multivariate study. Am J Pathol 150, 1997–2007 (1997).

38. Tanaka, K., et al. Prospective evaluation of Ki-67 system in histological grading of soft tissue sarcomas in the Japan Clinical Oncology Group Study JCOG0304. World J Surg Oncol 14, 110 (2016).

39. Woll, P.J., et al. Adjuvant chemotherapy with doxorubicin, ifosfamide, and lenograstim for resected soft-tissue sarcoma (EORTC 62931): a multicentre randomised controlled trial. Lancet Oncol 13, 1045–1054 (2012).

40. Cassinelli, G. The roots of modern oncology: from discovery of new antitumor anthracyclines to their clinical use. Tumori 2016, 226–235 (2016).

41. Kciuk, M., et al. Doxorubicin-An Agent with Multiple Mechanisms of Anticancer Activity. Cells 12(2023).

42. Tacar, O., Sriamornsak, P. & Dass, C.R. Doxorubicin: an update on anticancer molecular action, toxicity and novel drug delivery systems. J Pharm Pharmacol 65, 157–170 (2013).

43. Oshi, M., et al. A Novel Three-Gene Score as a Predictive Biomarker for Pathologically Complete Response after Neoadjuvant Chemotherapy in Triple-Negative Breast Cancer. Cancers (Basel) 13(2021).

44. Spalek, M.J., et al. Neoadjuvant Treatment Options in Soft Tissue Sarcomas. Cancers (Basel) 12(2020).

45. Alnoumas, L., et al. Evaluation of the role of KPNA2 mutations in breast cancer prognosis using bioinformatics datasets. BMC Cancer 22, 874 (2022).

46. Alshareeda, A.T., et al. KPNA2 is a nuclear export protein that contributes to aberrant localisation of key proteins and poor prognosis of breast cancer. Br J Cancer 112, 1929–1937 (2015).

47. Sun, Y., et al. Oncogenic role of karyopherin alpha2 (KPNA2) in human tumors: A pan-cancer analysis. Comput Biol Med 139, 104955 (2021).

48. Zhou, L.N., et al. Prognostic value of increased KPNA2 expression in some solid tumors: A systematic review and meta-analysis. Oncotarget 8, 303–314 (2017).

49. Xiang, S., et al. E2F1 and E2F7 differentially regulate KPNA2 to promote the development of gallbladder cancer. Oncogene 38, 1269–1281 (2019).

50. Yang, Y., et al. Silencing of karyopherin alpha2 inhibits cell growth and survival in human hepatocellular carcinoma. Oncotarget 8, 36289–36304 (2017).

51. Ma, A., et al. USP1 inhibition destabilizes KPNA2 and suppresses breast cancer metastasis. Oncogene 38, 2405–2419 (2019).

52. Sultana, A. & McMonagle, T. Pimozide for schizophrenia or related psychoses. Cochrane Database Syst Rev, CD001949 (2000).

53. Yu, Z., et al. USP1-UAF1 deubiquitinase complex stabilizes TBK1 and enhances antiviral responses. J Exp Med 214, 3553–3563 (2017).

54. Deutsch, E.W., et al. The ProteomeXchange consortium in 2020: enabling ‘big data’ approaches in proteomics. Nucleic Acids Res 48, D1145–D1152 (2020).

55. Perez-Riverol, Y., et al. The PRIDE database resources in 2022: a hub for mass spectrometry-based proteomics evidences. Nucleic Acids Res 50, D543–D552 (2022).

