## Supplemental Table 3 for "Proteomic characterisation of Sarculator nomogram-defined risk groups in soft tissue sarcomas of the extremities and trunk wall"

| **DepMap ID** | **Cell line** | **Histological subtype** |
| --- | --- | --- |
| ACH-001274 | SW982 | Synovial sarcoma |
| ACH-001275 | SYO1 | Synovial sarcoma |
| ACH-001277 | YAMATO | Synovial sarcoma |
| ACH-001280 | SCS214 | Synovial sarcoma |
| ACH-001322 | CME1 | Synovial sarcoma |
| ACH-001433 | CCLFPEDS0008T | Epithelioid sarcoma |
| ACH-001702 | VAESBJ | Epithelioid sarcoma |
| ACH-001791 | LPS6 | Liposarcoma |
| ACH-001793 | LPS27 | Liposarcoma |
| ACH-001799 | LPS141 | Liposarcoma |
| ACH-001804 | LPS510 | Liposarcoma |
| ACH-001802 | LPS853 | Liposarcoma |
| ACH-002785 | NCCLMS1C1 | Leiomyosarcoma |
| ACH-000145 | SKLMS1 | Leiomyosarcoma |
| ACH-000505 | RKN | Leiomyosarcoma |
| ACH-000939 | SKUT1 | Leiomyosarcoma |

Table S3. List of sarcoma cell lines form the Cancer Cell Line Encyclopaedia (CCLE) database used in this study
