## Supplemental Table 5 for "Proteomic characterisation of Sarculator nomogram-defined risk groups in soft tissue sarcomas of the extremities and trunk wall"

**Table S5** Univariable Cox regression analysis with Wald test assessing overall survival (OS) for patients stratified based on the sarculator nomogram risk groups and median expression of sarcoma proteomic module 6 (SPM6). HR=hazard ratio; CI= Confidence interval.

|  | **HR 95% CI p value** |
| --- | --- |
| **Sarculator nomogram**  3^rd^ vs 1^st^ quartile | 2.50 1.77-3.52 <0.0001 |
| **SPM6**  3^rd^ vs 1^st^ quartile | 1.43 0.94-2.19 0.2422 |
