## Supplemental Table 6 for "Proteomic characterisation of Sarculator nomogram-defined risk groups in soft tissue sarcomas of the extremities and trunk wall"

**Table S6** Multivariable Cox regression analysis with two-sided Wald test assessing interaction between sarcoma proteomic module 6 (SPM6) signature and tumour size. HR=hazard ratio; CI= Confidence interval.

|  | **HR 95% CI** |
| --- | --- |
| **Tumour size 1^st^ quartile (5.35 cm)**  **SPM6** | 1.504 0.77-3.09 |
| **Tumour size 2^nd^ quartile (8 cm)**  **SPM6** | 0.92 0.55-1.54 |
| **Tumour size 3^rd^ quartile (11 cm)**  **SPM6** | 0.87 0.48-1.58 |

Wald test p-value (p): interaction tumour size-SPM6 p=0.038; SPM6: linear term p=0.034; non-linear term p=0.008; tumour size: linear term p<0.0001; non-linear term p=0.07.
